# MET variants with activating N-lobe mutations identified in hereditary papillary renal cell carcinomas still require ligand stimulation

**DOI:** 10.1101/2023.11.03.565283

**Authors:** Célia Guérin, Audrey Vinchent, Marie Fernandes, Isabelle Damour, Agathe Laratte, Rémi Tellier, Gabriella O. Estevam, Jean-Pascal Meneboo, Céline Villenet, Clotilde Descarpentries, James S. Fraser, Martin Figeac, Alexis B Cortot, Etienne Rouleau, David Tulasne

## Abstract

In hereditary papillary renal cell carcinoma (HPRCC), the hepatocyte growth factor receptor (MET) receptor tyrosine kinase (RTK) mutations recorded to date are located in the kinase domain and lead to constitutive MET activation. This contrasts with MET mutations identified in non-small cell lung cancer (NSCLC), which lead to exon 14 skipping and deletion of a regulatory domain: in this latter case, the mutated receptor still requires ligand stimulation. Sequencing of MET in samples from 158 HPRCC and 2808 NSCLC patients revealed ten uncharacterized mutations. Four of these, all found in HPRCC and leading to amino acid substitutions in the N-lobe of the MET kinase, proved able to induce cell transformation, which was further enhanced by hepatocyte growth factor (HGF) stimulation: His1086Leu, Ile1102Thr, Leu1130Ser and Cis1125Gly. Similar to the variant resulting in MET exon 14 skipping, the two N-lobe MET variants His1086Leu and Ile1102Thr were found to require stimulation by HGF in order to strongly activate downstream signaling pathways and epithelial cell motility. The Ile1102Thr mutation also displayed transforming potential, promoting tumor growth in a xenograft model. In addition, the N-lobe-mutated MET variants were found to trigger a common HGF-stimulation-dependent transcriptional program, consistent with an observed increase in cell motility and invasion. Altogether, this functional characterization revealed that N-lobe variants still require ligand stimulation, in contrast to other RTK variants. This suggests that HGF expression in the tumor microenvironment is important for tumor growth. The sensitivity of these variants to MET inhibitors opens the way for use of targeted therapies for patients harboring the corresponding mutations.

## Introduction

Receptor tyrosine kinases (RTKs) play an important role in different cell processes, including cell proliferation, motility, differentiation, and metabolism[1,2]. Fifty-eight RTKs have been described in humans, all sharing a similar protein structure with an extracellular ligand-binding domain, a single transmembrane domain, and an intracellular domain containing a regulatory juxtamembrane domain, a tyrosine kinase domain, and a carboxy-terminal tail[3]. Importantly, dysregulations of RTK signaling are associated with the development of many cancers. With the advent of next-generation sequencing for molecular diagnosis, exhaustive mutational landscapes have emerged for almost all types of tumors[4]. In particular, several types of alteration have been found in genes encoding RTKs. As a representative example, epidermal growth factor receptor (EGFR)[5] displays several amino-acid substitutions and deletions within its kinase domain, leading to its constitutive ligand-independent activation, notably in lung cancer[6].

Over the last two decades, RTKs have emerged as the main targets for targeted therapies, with the development of many small-molecule tyrosine kinase inhibitors (TKIs) approved for use against numerous cancers [7]. All the currently FDA-approved TKIs act as ATP competitors, able to associate with the ATP-binding pocket of the kinase domain in its active form (type I inhibitors) or inactive form (type II inhibitors) [8].

MET, another attractive RTK target for cancer treatment, is found predominantly in epithelial cells and is activated by its high-affinity ligand, the hepatocyte growth factor (HGF)[9]. MET is activated through homo-dimerization upon HGF binding to its extracellular domain. Then, each tyrosine kinase transphosphorylates tyrosine residues Y1234 and Y1235 within the catalytic loops, promoting further kinase activity. This activation leads to phosphorylation of other tyrosine residues, including Y1349 and Y1356 in the C-terminal tail, involved in recruitment of signaling proteins leading to activation of downstream signaling and of cellular responses including proliferation, motility, survival, and invasion[10]. In mammals, the HGF/MET pair is essential to embryonic development[9] and to tissue regeneration in adults[11–13].

Several genomic events lead to MET activation and oncogene-driven tumors[14]. These include MET gene amplification (mainly observed in stomach and lung cancer), causing overexpression and ligand-independent activation of MET[15–21], alterations (mostly observed in glioblastoma) causing HGF overexpression leading to aberrant MET activation through establishment of an autocrine loop[22,23], and chromosomal rearrangements leading to MET fusion with another gene (observed in lung adenocarcinoma and renal cell carcinoma), inducing either constitutive activation or overexpression of MET[24]. Many MET mutations have also been described in hereditary papillary renal cell carcinoma (HPRCC)[25]and non-small cell lung cancer (NSCLC), with for each cancer specific types of mutations.

Papillary renal cell carcinoma (PRCC), representing 15-20% of renal cell carcinomas (RCCs), includes two main histological subtypes: hereditary PRCC (HPRCC), known as type 1, and non-hereditary PRCC, known as type II[26–28]. MET-activating mutations were initially described in HPRCC, the hereditary character of these alterations providing first evidence of an involvement of MET in tumorigenesis[25,29,30]. Since then, many MET-activating mutations, located in the kinase domain, have been identified in non-hereditary papillary renal cancers[31] and found to induce constitutive MET activation leading to cell transformation and tumor growth in experimental models. In a phase II clinical trial evaluating foretinib, a multikinase inhibitor which notably targets MET, half of the ten PRCC patients harboring a germline MET mutation displayed a positive response [32]. As this suggests the possibility of treating such patients with MET TKIs, several MET inhibitors are currently under clinical evaluation (https://www.clinicaltrials.gov/search?term=HGFR&cond=Renal%20Cell%20Carcinoma).

In non-small cell lung cancer (NSCLC), about 3% of patients display a wide variety of mutations and deletions, all leading to skipping of MET exon 14 (METex14Del) [16,33–35], encoding the negative regulatory juxtamembrane domain of the receptor. Yet unlike the mutated RTK variants classically described and despite its oncogenic character, the MET variant resulting from exon 14 skipping still requires stimulation by HGF for its oncogenicity[36,37]. Several MET TKIs have recently been approved for treating patients harboring MET exon 14 skipping mutations[38–41]. Yet systematically, resistances to these targeted therapies have been observed. Several mechanisms for this have been described, mainly activation of other RTKs or downstream signaling effectors[42,43] and on-target mutations located in the MET kinase domain[44–52]. Interestingly, some of the MET mutations involved in resistance have already been described in renal cancer[25,53–55].

Molecular diagnosis, based on genomic sequencing of oncogene drivers including MET, has led to the recent identification of novel somatic variants, whose functional involvement and potential sensitivity to TKIs are seldom known. Thus, the challenge is no longer to identify RTK variants but to characterize them functionally. Here we have profiled, in samples from a cohort of 2808 NSCLC and 158 HPRC patients, the molecular alterations affecting the MET receptor. We have performed a full functional characterization of the corresponding MET variants, focusing on cell transformation, induction of signaling pathways and biological responses, transcriptional programs, and experimental tumor growth to improve our understanding of these variants and their targeting.

## Materials and methods

### Patient cohorts

One hundred and fifty-eight patients with PRCC1 from the Gustave Roussy Institute were screened for pathogenic germline MET variants from 2001 to 2019 as previously described [56]. MET genetic screening was performed on DNA extracted from peripheral blood leukocytes. Next-generation sequencing (NGS) was performed and the genetic variants detected were confirmed by Sanger Sequencing or multiplex ligation probe-dependent amplification (MLPA SALSA MLPA Probemix P308 MET; MRC Holland). For NSCLC, 2808 patient samples from Lille University Hospital were analyzed with the CLAPv1 NGS panel [57]. A detailed statement for personal data processing has been made to the Lille University Hospital Data Protection Officer, under the n°273. For GDPR, the patients have been informed using medical letters (with mention of possible data re-using and possible opposition), and/or a specific information letter about the study. The possible oppositions have been researched into medical files and on Hospital Information System. A Privacy Impact Assessment has been performed before any data treatment. All the data have been anonymized before collection, storage and treatment. Both PRCC and NSCLC methodologies are conformed to the standards set by the Declaration of Helsinki and were approved by the local ethics committee.

### Cell line maintenance

NIH3T3 (NIH/3T3) (RRID:CVCL_0594) cells were cultured in DMEM, high glucose, Glutamax (Invitrogen, Carlsbad, CA) supplemented with 10% bovine calf serum (Life technologies, New Zealand) and 1% Zell-shield antibiotics (Cliniscience, Germany). The MCF-7 (RRID:CVCL_0031) parental cell line and derivatives thereof were maintained in DMEM, high glucose, Glutamax supplemented with 10% heat-inactivated FBS (Sigma, St. Louis, Missouri), 1% nonessential amino acids (Gibco, Invitrogen, Scotland), and 1% Zell-shield antibiotics (Minerva Biolabs, Berlin, DE). Cells were cultivated at 37°C under a humidified controlled atmosphere of 5% CO2 in air. Mycoplasma tests were routinely performed with the MycoAlert mycoplasma detection kit (Lonza, Basel, Switzerland). Cell lines were obtained at ATCC and MCF-7 cells have been authentified in the past three years by genetic characteristics determined by PCR-single-locus-technology based on 16 independent PCR-systems D8S1179, D21S11, D7S820, CSF1PO, D3S1358, TH01, D13S317, D16S539, D2S1338, AMEL, D5S818, FGA, D19S433, vWA, TPOX and D18S51.

### Plasmid engineering

MET mutations causing MET p.(Val37Ala), MET p.(Arg426Pro), MET p.(Ser1018Pro), MET p.(Gly1056Glu), Ile1102Thr, MET p.(His1086Leu), MET p.(Leu1130Ser), MET p.(Cys1125Gly), and MET p.(Met1268Thr) were created as previously described [58] with the QuickChange site-directed mutagenesis system (Agilent Technologies, Santa Clara, CA) on the pRS2-MET vector encoding wild-type MET cDNA. Each MET cDNA was then subcloned with EcoR1 into a pCAGGS expression vector. All MET-encoding sequences were completely verified by Sanger sequencing.

### Stable expression of wild-type and mutant MET in the MCF-7 cell line

MCF-7 cells were transfected by Lipofectamine 2000 (Invitrogen, Netherlands) with the pCAGGS vector expressing MET WT or the mutant MET cDNA and the pSV2 plasmid expressingg the neomycin resistance gene (ratio 10:1). The next day, the cells were treated with G418 (ThermoFisher Scientific, Waltham, MA) at 800 μg/ml during 21 days. Clones were collected on filter paper and MET expression was assesses by western blotting, immunofluorescence, and qPCR.

### Stable expression of wild-type and mutant MET in the NIH3T3 cell line

NIH3T3 (NIH/3T3) cells (250,000 cells in a six-well plate) were transfected with Lipofectamine 2000 (Invitrogen, Netherlands) (per well, 2 µg DNA/4 µl Lipofectamine 2000 in 400 µl Opti-MEM medium; Thermo Fisher Scientific). The next day, the cells were transferred to a 100-mm plate and, after reaching confluence two days later, cultured in 5% calf serum. After fifteen days, clones were isolated on filter paper. Western blot and qPCR analyses confirmed MET expression.

### NIH3T3 transfection and transformation

*In vitro* transforming ability assays are based on the capacity of NIH3T3 cells to form foci when transfected with a mutant MET. The mutants tested were MET p.(Val37Ala), MET p.(Arg426Pro), MET p.(Ser1018Pro), MET p.(Gly1056Glu), MET p.(Ile1102Thr), MET p.(His1086Leu) and MET p.(Cys1125Gly). Two hundred and fifty thousand NIH3T3 cells plated in a 6-well plate were transfected in the presence of Lipofectamine 2000 (Invitrogen) (per well 2 µg DNA/4 µl Lipofectamine 2000 in 400 µl Opti-MEM medium; Thermo Fisher Scientific). The next day, the cells were transferred to a 100-mm plate and, after reaching confluence two days later, cultured in 5% calf serum in the presence or not of HGF (10ng/ml). Fifteen days later, the cells were fixed in fast green/methanol and stained with Carazzi/eosin. Results were compared with those obtained with NIH3T3 cells transfected with a construct encoding the strongly activating MET mutant MET p.(Met1268Thr) or wild-type MET.

### TKIs and cytokines

HGF was purchased from Miltenyi Biotec (Bergisch Gladbach, Germany). The MET inhibitors crizotinib, merestinib, and capmatinib were from Selleck Chemical, USA. HGF blocking antibody Rilotumumab (HY-P99217) and human IgG2 Kappa, isotype control (HYP99002) were purchased from MedChemExpress (New Jersey, USA).

### Western blot analyses

Cells were plated on 6-well plates (250,000 cells per well) in complete medium. The next day, the cells were serum-starved overnight and treated with HGF in serum-free medium before lysis. They were washed in PBS and suspended in lysis buffer (20 mM Tris-HCl (pH 7.4), 50 mM NaCl, 5 mM EDTA, 1% (v/v) Triton X-100, 0.02% (w/v) sodium azide, and water) containing freshly added protease and phosphatase inhibitors (1% Aprotinib, 1mM PMSF, 1mM leupeptin, 1mM Na_3_VO_4_, and β glycerophosphate). Lysates were clarified by centrifugation at 4°C and the protein concentration was determined by BCA protein assay (Thermofisher). Samples were resolved on a precast 4%–12% gradient gel (Thermo Fisher Scientific) and transferred to Immobilon-P (Merck Millipore) membranes.

The following antibodies were used: MET (Cell Signaling #3148), MET-Phospho (Cell Signaling #3126), AKT (Cell Signaling #2920), AKT-Phospho (Cell Signaling #4060), ERK1/2 (Santa Cruz #Sc-154), ERK1/2-Phospho (Cell Signaling #9101), GAPDH (Santa Cruz #Sc-32233/6c5). Peroxidase-coupled secondary antibodies were purchased from Jackson ImmunoResearch Laboratories (anti-mouse 115-0350146; anti-rabbit 711-0350152). Proteins were detected with SuperSignal™ West Dura (Thermofisher) and SuperSignal™ West Femto (Thermofisher). Luminescence was captured by digital imaging with a cooled charge-coupled device camera (ChemiDoc - BioRad) and analyzed with ImageLab software.

### RT-qPCR

RT-qPCR were performed as previously described [58]. Briefly, total RNA was extracted with the Nucleospin RNA/Protein kit (Macherey-Nagel, Düren, DE) according to the manufacturer’s instructions. cDNA was obtained by reverse transcription with random hexamers (Applied Biosystems, Foster city, CA). Levels of MET and target gene mRNAs were evaluated by real-time RT-PCR with Fast SYBR Green mix (Applied Biosystems). Relative gene expression levels were calculated using the 2-ΔCt method. The level of each transcript was normalized to GAPDH. The primers used are listed in Table 1.

**Table 1:** RT-qPCR primers.

### Migration assays

Migration assays were performed as previously described [58]. Briefly, for IncuCyte® Scratch Wound 96-Well Real-Time Cell Migration assays (wound healing), cells were seeded at 30,000 cells/well in a 96-well plate. DMSO, HGF (30 ng/ml), and the MET TKI inhibitor capmatinib (1 μM), alone or together, were added, and cells were placed in the IncuCyte Zoom live cell imaging system (Essen BioScience) and Incucyte SX5 live cell imaging system (Sartorius). Pictures were taken every 3 h for 96 h. Custom algorithms of the IncuCyte™ software package calculated wound healing.

### Transcriptomic analysis

Cells were plated on Petri dishes in complete medium (750,000 cells/dish). The next day they were serum-starved for 2 h and treated or not with HGF in serum-free medium for 24 h. Total RNA was extracted with the Nucleospin RNA kit (Macherey-Nagel) according to the manufacturer’s instructions. Genomic DNA traces were removed by DNAse I treatment. HGF stimulation and RNA extraction were performed four times at one-week intervals. Total RNA yield and quality were evaluated on the Nanodrop 2000C system (ThermoScientific) and further assessed on the Agilent 2100 bioanalyzer (Agilent Technologies).

#### Construction of sequencing libraries

Starting from 4 ul total RNA, we added 1 ul ERCC spike-in control. Library generation was then initiated by oligo dT priming, from 200ng of total RNA. The primer already contained Illumina-compatible linker sequences (Read 2). After first strand synthesis, the RNA was degraded and random-primed second strand synthesis was initiated by DNA polymerase. The random primer also contained 5’ Illumina-compatible linker sequences (Read 1). At this step Unique Molecular Identifiers (UMIs) were introduced, allowing elimination of PCR duplicates during analysis. After obtaining the double-stranded cDNA library, the library was purified with magnetics beads and amplified. During library amplification, the barcodes and sequences required for cluster generation (index i7 in 3’ and index i5 in 5’) were introduced thanks to Illumina-compatible linker sequences. The number of cycle of amplification was 14. The final library was purified and deposited on a high-sensitivity DNA chip to be checked on an Agilent 2100 bioanalyzer. The library concentration and size distribution were checked.

#### Sequencing

The libraries were pooled equimolarly and the final pool was also checked on an Agilent 2100 bioanalyzer and sequenced on NovaSeq 6000 (Illumina) with 100 chemistry cycles.

#### Analysis of transcriptomic data

To eliminate poor quality regions and poly(A) from the reads, we used the fastp program. We used a quality score threshold of 20 and removed reads shorter than 25 pb. Read alignments were performed with the STAR program, using the reference human genome (GRCh38) and reference gene annotations (Ensembl). The UMI (Unique Molecular Index) allowed reducing errors and quantitative PCR bias with fastp and umi-tools. On the basis of read alignments, we counted numbers of molecules per gene with FeatureCount. Other programs, such as as qualimap, fastp, FastQC and MultiQC, were performed for read quality control. Differential gene expression analysis of RNA-seq data was perfomed with the R/Bioconductor package DESeq2. The cut-off for differentially expressed genes was fold change > 1.5 and p-value padj (BH) < 0.05. This method have also been described in our previous study [58].

### In vivo tumor xenografts

The project (19253-201903191709966 v1) received ethical approval by French Committee on Animal Experimentation and the Ministry of Education and Research. Mice were housed in standard conditions, with a 12-hour light/dark cycle, controlled temperature (20-24°C), and humidity (40-60%). They were kept in ventilated cages with ad libitum access to food and water. Bedding was changed regularly to maintain hygiene. Xenografts were performed on CB-17/lcr-*Prkdc^scid/scid^*/Rj (SCID) mice from the Institut Pasteur de Lille, France (Agreement B59-350009). NIH3T3 cells (2x10^6^) in PBS were injected subcutaneously into both flanks of 9 weeks mice with an equal repartition between males and females. Tumors were palpated and measured with calipers at least twice a week until they reached 100 mm^3^ in volume. The tumor volume (V) was calculated with the formula V=0.5 x (LxW^2^). Animals were culled before the tumors reached 1500 mm^3^ or if limit points were reached.

### Immunofluorescence

After the tissue sections were deparaffinized heat-induced epitope retrieval was performed. The slides were incubated with primary antibodies against MET (AF276 R&D systems at 0.2μg/ml) and anti-goat antibody conjugated with Alexa 488 (A11055, Invitrogen). Slides were stained with Hoechst mounted with Moviol mounting medium. For fluorescence microscopy, slides were observed in oil immersion with a Zeiss AxioImager Z1 ApoTome microscope.

### Statistics

All results are expressed as means±s.e.m. of at least three independent measurements. According to their distribution, quantitative variables were compared with a t-test or a two-way ANOVA. T-tests were performed at each time point for tumor growth analyses in subcutaneous xenograft models. All statistical testing was conducted at the 2-tailed α level of 0.05. Data were analyzed with GraphPad Prism software version 10.

## Results

### Focus formation induced by N-lobe MET mutations is enhanced by HGF

A set of 158 samples from PRCC type I (HPRCC) patients was analyzed by NGS for the presence of germline MET mutations. Eight germline point mutations inducing amino-acid substitutions were identified as undescribed pathogenic or likely pathogenic MET mutations[59]: MET p.(Val37Ala) and MET p.(Arg426Pro) affecting the SEMA domain, MET p.(Ser1018Pro) and MET p.(Gly1056Glu) affecting the juxtamembrane domain, and MET p.(His1086Leu), MET p.(Cys1125Gly), MET p.(Ile1102Thr), and MET p.(Leu1130Ser) affecting the N-lobe region of the kinase domain (Table 2 and Figure 1A and B). All these variants were annotated from the MET transcript variant 1 NM_001127500.1. We have recently started characterizing the His1086Leu, Cys1125Gly, Ile1102Thr, and Leu1130Ser variants and have found that without HGF stimulation, they display weak oncogenicity [59]. In parallel, a set of 2808 NSCLC patient samples were analyzed with the CLAPv1 NGS panel[57]. Two-point mutations inducing amino-acid substitutions were detected within the kinase domain: His1097Arg and Asp1249Glu (Table 2). To investigate the oncogenicity of these novel MET mutations, we first constructed three control vectors: one expressing the wild-type MET cDNA, one expressing a MET cDNA containing the known MET p.(Met1268Thr) mutation, which induces ligand-independent MET activation, and one expressing a cDNA carrying the activating MET p.(Met ex14) mutation (the deletion due to exon 14 skipping)(METex14Del) that requires ligand stimulation[36,37]. We also constructed expression vectors encoding the MET variants corresponding to the various novel mutations identified in HPRCC and NSCLC tumors. In what follows, for the sake of simplicity, we use for each mutated form of the protein the name of the corresponding genomic mutation.

**Figure 1:**
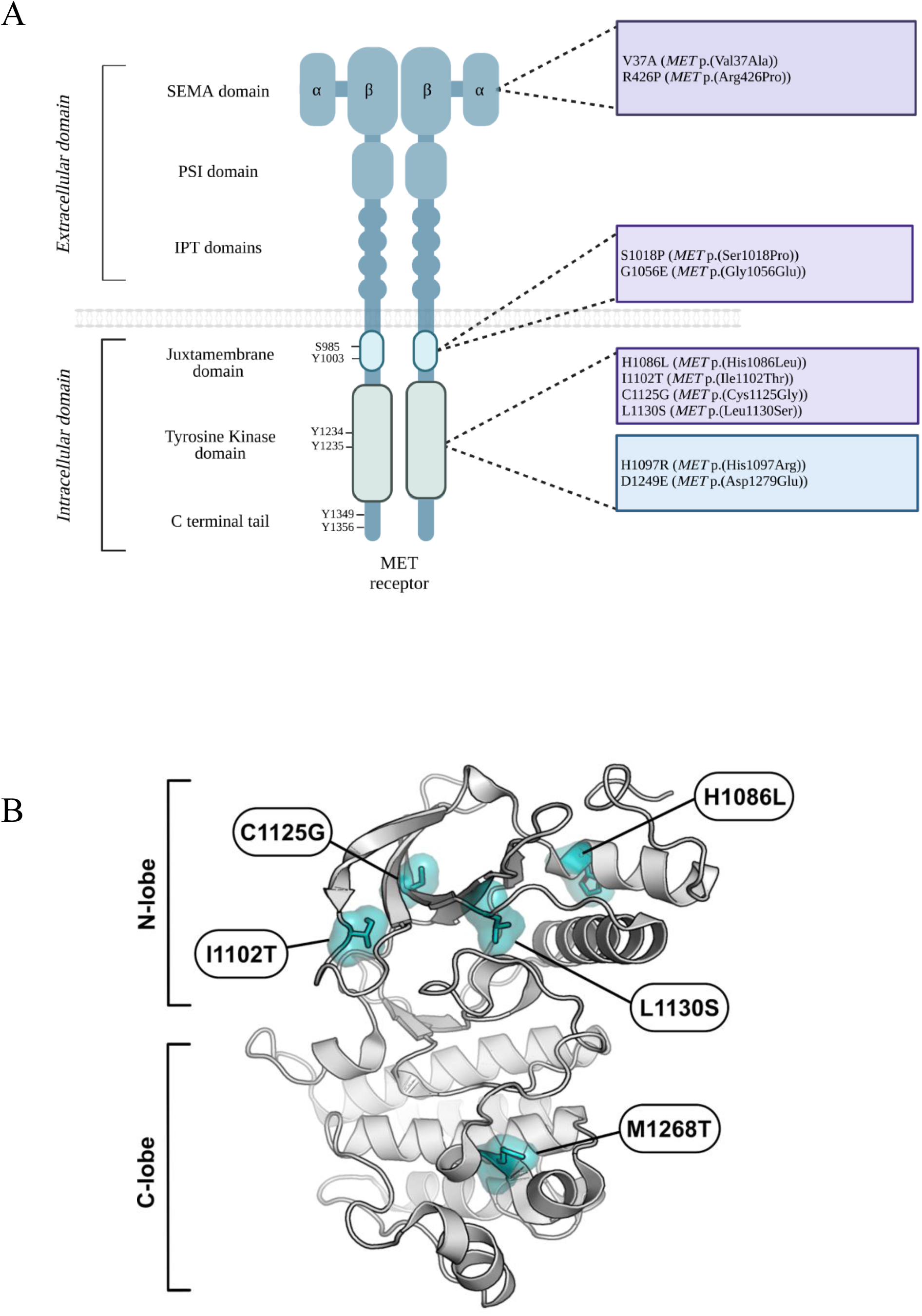
Schematic representation of MET variants. (A) Representation of novel MET mutations in the SEMA domain, the juxtamembrane domain, and the tyrosine kinase domain, identified in HPRCC (in purple) and NSCLC (in blue). (B) Crystal structure of the human MET kinase domain (PDB 2G15). Highlighted are, on the one hand, the documented activating mutation M1268T in the C-lobe of the kinase and, on the other hand, the novel N-lobe mutations characterized in this study, analyzed through PyMOL.

**Table 2:** Description of MET variants uncharacterized localized in SEMA, juxtamembraine and kinase domain.

We first performed in vitro transformation assays on the NIH3T3 cell line, with and without HGF stimulation (Figure 2). The Met1268Thr mutant used as positive control induced, as expected, abundant focus formation, independently of the presence or absence of HGF. The kinase mutations His1097Arg and Asp1249Glu, found in NSCLC tumors, failed to induce focus formation whether HGF was present or not, which suggests that they have no transforming ability. The same was true of the following variants found in HPRCC: Val37Ala and Arg426Pro (SEMA domain), Ser1018Pro and Gly1056Glu (juxtamembrane domain) (Figure 2A, B and C, D). In contrast, Leu1130Ser, Cys1125Gly, Ile1102Thr, and His1086Leu, found in HPRC patients, induced minor focus formation in the absence of HGF but enhanced focus formation upon HGF stimulation (Figure 2E, F and G, H). Expression levels of all MET variants were assessed by western blotting (Figure 2A, C, E, and G). It is worth noticing that all MET construct display phosphorylation on the tyrosine residues of the kinase domain without HGF stimulation due to strong expression induced by transient transfection.

**Figure 2:**
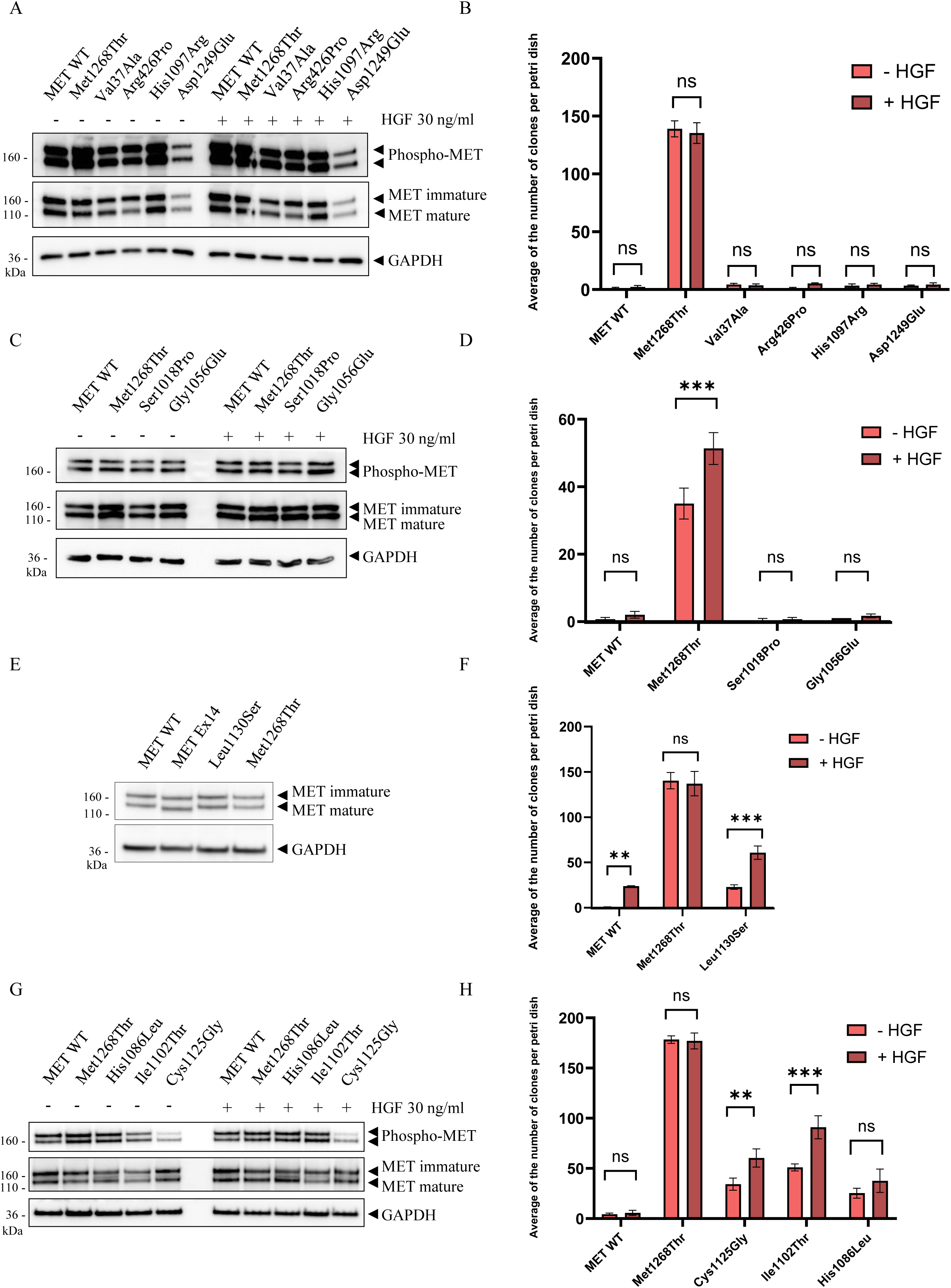
In vitro transforming ability assays performed on cells expressing different mutated MET cDNAs in the absence and presence of exogenous HGF. Western blot showing transient MET expression (A, C, E, G) and diagrams showing the average number of clones per Petri dish (B, D, F, H) of NIH3T3 cells transfected with a vector encoding either wild-type (WT) MET, MET p.(Met1268Thr) (positive control), or a novel MET variant. (A and B) Results for MET p.(Val37Ala), MET p.(Arg426Pro), MET p.(His1097Arg) and MET p.(Asp1249Glu). (C and D) Results for MET p.(Ser1018Pro) and MET p.(Gly1056Glu). (E and F) Results for MET p.(Leu1130Ser). (G and H) Results for MET p.(Cys1125Gly), Ile1102Thr and MET p.(His1086Leu). For Western botting, cells were stimulated 30min by 30ng/ml HGF. For foci formation assay, cells were treated all along the culture by 10ng/ml HGF. n=3; mean ± SEM ; representative of three independent experiments. Statistical analysis by two-way ANOVA, ** p-value <0.0021; *** p-value <0.0002.

These novel MET-activating mutations were mapped on the structure of the kinase domain (Figure 1B). They are located in the N-lobe of this domain, within the ATP-binding loop (also called the P-loop). More precisely, the His1086Leu substitution is located in the ɑJM-helix (aJuxtamembrane helix) and the Ile1102Thr substitution is close to the ATP-binding loop.

On the basis of these findings, we pursued our functional analysis of the Leu1130Ser, Cys1125Gly, Ile1102Thr and His1086Leu kinase variants corresponding to the genomic mutations found in HPRCC tumors. We first evaluated their sensitivity to three MET TKIs: the type I TKIs crizotinib and capmatinib and the type II TKI merestinib (Figure 3). MET phosphorylation of all four variants was inhibited by all three TKIs, which suggests potential sensitivity to MET-targeted therapy. Interestingly, alignement of MET kinase domain with other kinases indicate that residues mutation in a similar position that MET Ile1102 induce resistance to ATP competitors in the CDK6, EGFR, ERK2, BRAF, and HER2 kinases [60], this was not the case for MET.

**Figure 3:**
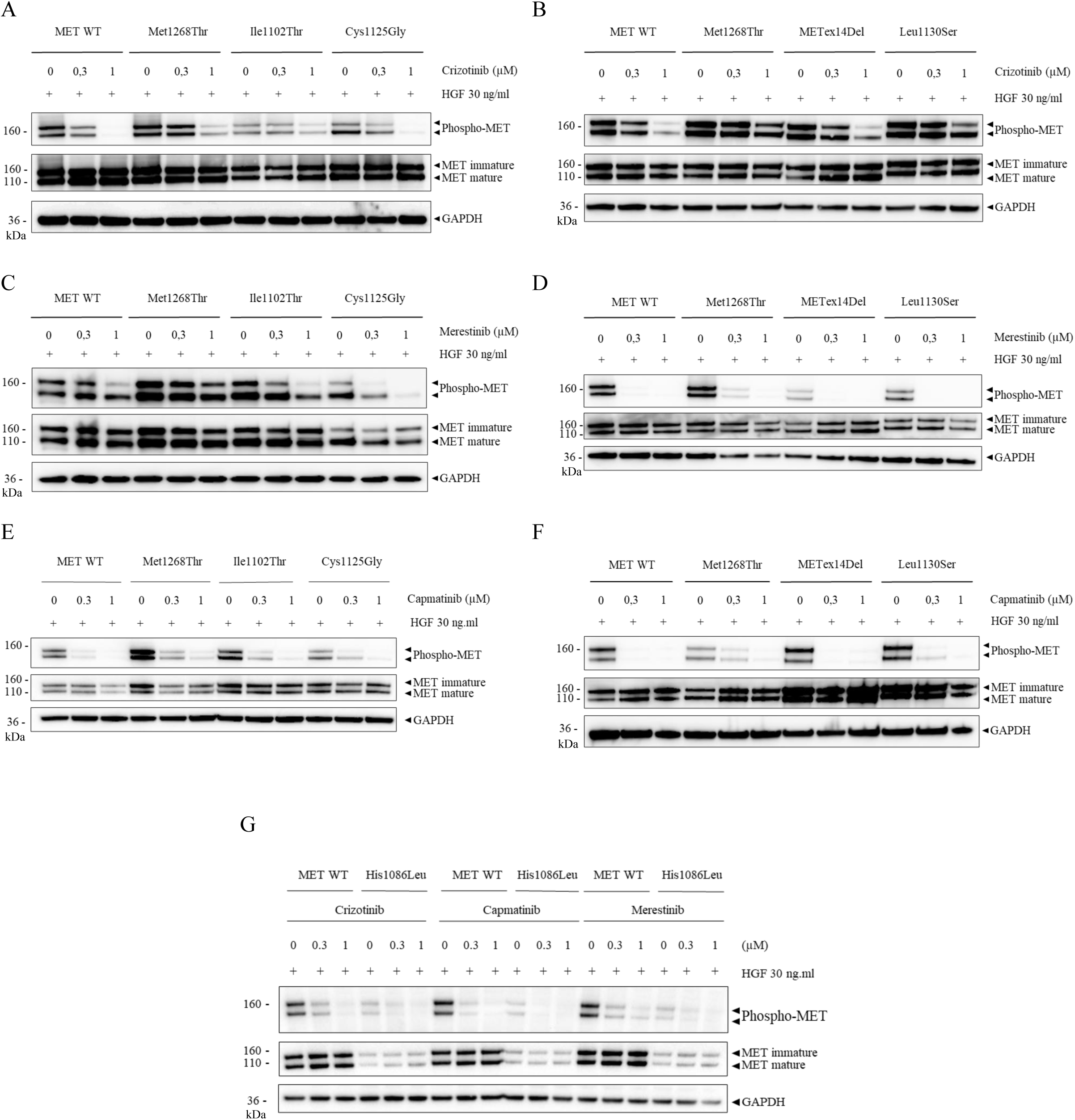
MET TKI sensitivity and inhibition of downstream signaling pathways in transiently tranfected NIH3T3 cells expressing different MET variants. Western blot showing the MET activity (phospho-MET and MET) upon TKI treatment (Tyrosine Kinase inhibitor crizotinib, merestinib, or capmatinib at the indicatd concentration for 1.5h), prior stimulation or not 30min by 30ng/ml HGF of NIH3T3 cells expressing either MET p.(His1086Leu) (H1086L), MET p.(Ile1102Thr) (I1102T), MET p.(Cys1125Gly) (C1125G), or MET p.(Leu1130Ser) (L1130S), as compared to cells expressing the wild-type MET and to control cells expressing MET p.(Met1268Thr) (M1268T) or METex14Del. GAPDH expression was studied as quality control to validate the experiment. (A and B) MET activity of the I1102T, C1125G and L1130S MET mutants after crizotinib treatment. (C and D) MET activity of the I1102T, C1125G, and L1130S MET mutants after merestinib treatment. (E and F) MET activity of the I1102T, C1125G, and L1130S MET mutants after capmatinib treatment. (G) MET activity of the H1086L MET mutant after crizotinib, merestinib or capmatinib treatment. These western blots are representative of three independent experiments.

### HGF stimulation enhances HGF-induced epithelial cell responses

In order to investigate and characterize the biological responses induced by MET in epithelial cells, MCF-7 epithelial cells of mammary origin were stably transfected with an expression vector encoding either the MET wild type or a mutated MET form. Advantages of this cell line are that it does not express MET endogenously[61]. In this model, we focus our attention on the two N-lobe mutations Ile1102Thr and His1086Leu and the METex14Del variant known to require ligand stimulation. MET expression was assessed by western blotting (Figure 4A) and RT-qPCR (Figure 4B). As expected, the parental MCF-7 cells showed no expression. Similar expression levels were observed for the MET variants.

**Figure 4:**
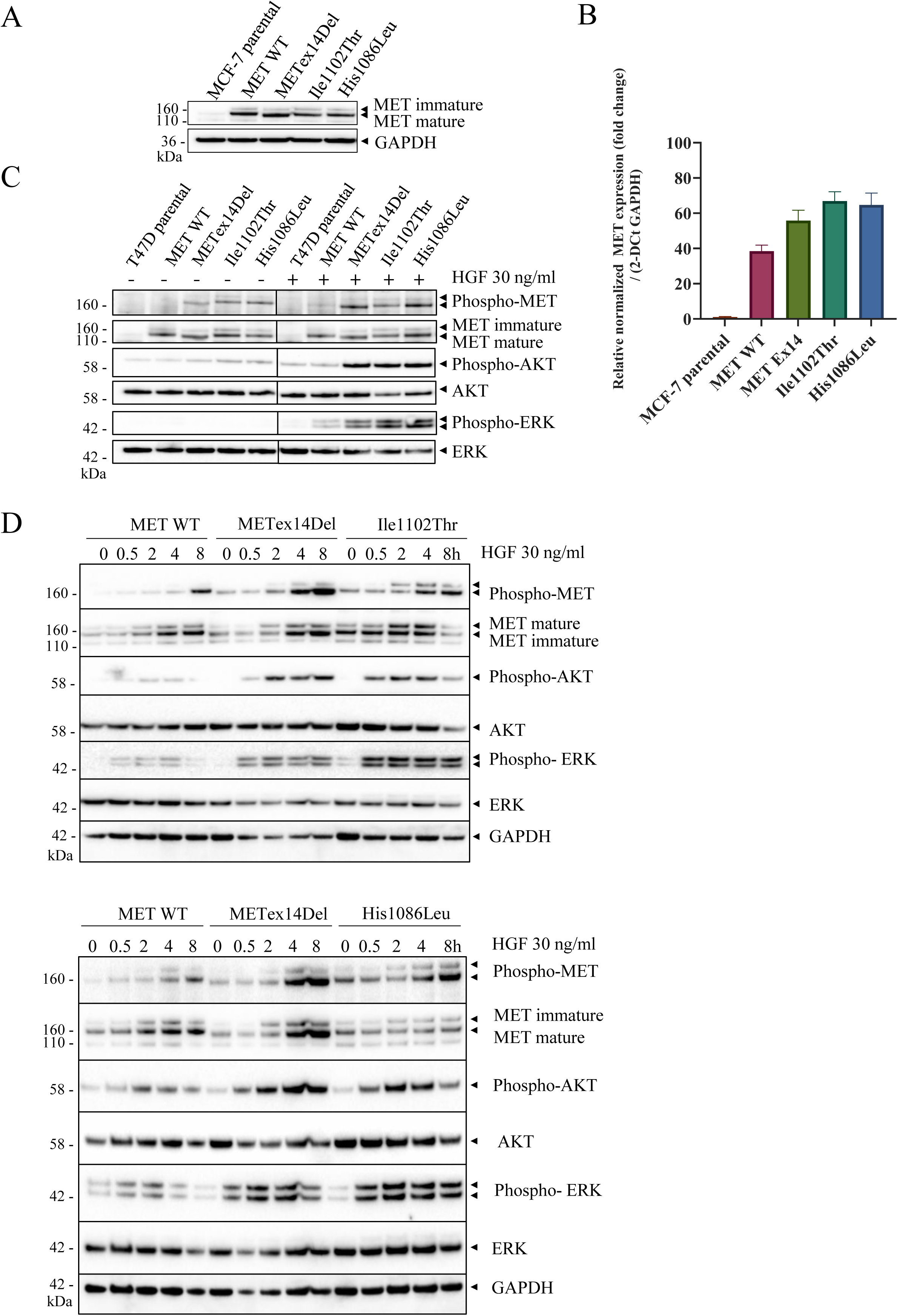
MET expression in MCF-7 cells as determined by western blotting and qPCR. Downstream signaling pathway activation in MCF-7 cells. (A) MCF-7 cells expressing wild-type (WT) or mutated MET were plated at 250,000 cells and lysed 24h later. MET levels were determined by western blotting with the indicated antibodies. GAPDH was used as loading control. (B) MET expression was also validated by RT-qPCR. 250,000 cells were seeded and 24h later, total RNA was extracted and complementary DNA (cDNA) was obtained with a reverse transcriptase kit. n=3; mean ± SEM. (C) MCF-7 cells were incubated overnight in serum-free medium and stimulated or not for 30 min with 30 ng/ml HGF prior to cell lysis. Levels and phosphorylation of MET, AKT, and ERK, were determined by western blotting with the indicated antibodies. ERK was used as loading control. (D) MCF-7 cells were incubated overnight in serum-free medium and stimulated or not for 30min, 2h, 4h or 8h with 30 ng/ml HGF prior to cell lysis. Levels and phosphorylation of MET, AKT, and ERK were determined by western blotting with the indicated antibodies. ERK and GAPDH were used as loading control. The western blots are representative of three independent experiments.

HGF stimulation during 30 minutes did induce ERK and AKT phosphorylation, which was more pronounced in cells synthesizing a mutated MET form than in ones synthesizing MET wild type (Figure 4C). Except for MET wild type, all the MET variants displayed basal MET phosphorylation in the absence of HGF stimulation. At this time of stimulation, phosphorylation was not further enhanced by HGF stimulation (Figure 4C). Time course experiments showed that MET expression and phosphorylation increased for all MET construct from 2h until 8h stimulation by HGF, which was more pronounced for METex14Del followed Ile1102Thr and His1086Leu. AKT phosphorylation induced by METex14Del was sustained for 8h, whereas with Ile1102Thr and His1086Leu, a decrease in AKT phosphorylation was observed at 8h (Figure 4D). ERK phosphorylation was sustained for all the mutated forms. The sustained AKT activation by METex14Del is consistent with previous studies indicating that loss of the juxtamembrane domain prevents downregulation of MET activity and signaling[62,63].

In wound healing experiments, all the cell lines displayed similar, low-level wound healing in the absence of HGF stimulation. Under HGF stimulation, parental cells showed no increase in wound healing, while MET wild-type cells displayed a minor increase (about 50% at 48 h). HGF stimulation induced a quicker wound healing response in cells expressing Ile1102Thr, His1086Leu, or METex14Del (about 90% at 48 h) (Figure 5A and B). These responses were abolished by treatment with the MET tyrosine kinase inhibitor crizotinib (Figure 5C) and by blocking antibody directed against HGF (Supplementary Figure S1). MET TKIs, including crizotinib, merestinib and capmatinib, were also able to strongly decrease, MET, ERK and AKT phosphorylation induced by HGF in MCF-7 cells expressing MET WT, METex14Del, Ile1102Ther and His1086Leu (Supplementary Figure S2A, B and C). Therefore, without HGF stimulation, cell motility was comparable for all the MCF-7 cell lines, but in response to HGF stimulation, cells expressing METex14Del, Ile1102Thr, or His1086Leu displayed drastically increased motility, which was sensitive to MET TKI and blocking antibody. They also showed increased intracellular signaling as compared to MET wild-type cells.

**Figure 5:**
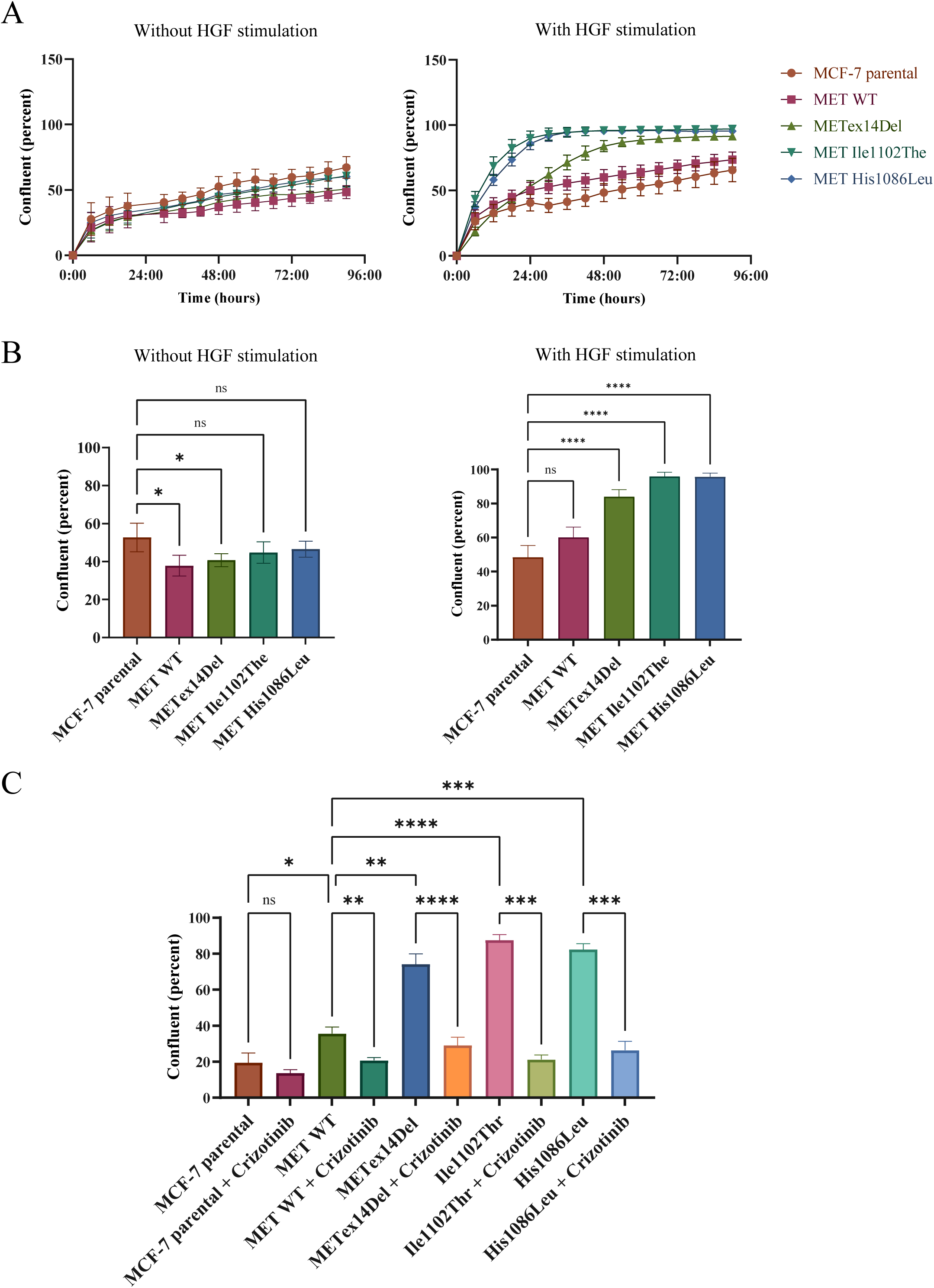
Migration prompted by HGF-stimulated MET mutants in MCF-7 cells. (A) MCF-7 cells were seeded at 30,000 cells per well in a 96-well plate. Twenty-four hours later, mitomycin C (10 µg/ml) was added for 2h to stop proliferation. Then a scratch wound was performed with the woundmaker tool and cells were stimulated or not with HGF at 10 ng/ml. Pictures were taken every 3 h for 96 h. Data are expressed as mean relative proliferation (n=3; mean ± SEM). (B) 48h points of wound healing are shown (n=3; mean ± SEM). (C) MCF-7 cells were seeded at 30,000 cells per well in a 96-well plate. Twenty-four hours later, mitomycin C (10 µg/ml) with or without crizotinib (1 µM) was added for 90 min to stop proliferation and inhibit MET. Then a scratch wound was performed with the woundmaker tool and cells were stimulated with HGF at 10 ng/ml. 30h points of wound healing are shown. Data are expressed as mean relative proliferation (n=3; mean ± SEM). Statistical analysis by one-way ANOVA, * p-value <0.0332; ** p-value <0.0021; *** p-value <0.0002; **** p-value <0.0001.

### Activating N-lobe mutations induce a specific transcriptional program promoting cell motility

To further characterize the responses induced by each MET variant upon HGF stimulation, the transcriptomes of the different MCF-7 cell lines were compared. Volcano plots (Figure 6A) present the genes differentially expressed according to whether the cells were stimulated with HGF or not (fold change >1.5 and adjusted p-value </= 0.5). In the parental cell line, HGF stimulation induced no modification of gene expression. In cells expressing the MET wild type, only 41 genes appeared differentially expressed. In contrast, HGF stimulation induced drastic modifications in gene expression in the cells expressing mutant MET cDNAs, with 4311 differentially expressed genes for METex14Del, 3327 for Ile1102Thr, and 2566 for His1086Leu. The list of the differentially regulated genes in response to HGF is shown in the Supplementary Table 1 A, B, C, D and E. In the absence of HGF stimulation, global unsupervised clustering of the gene expression profiles (Figure 6B) showed the MCF-7 parental METex14Del, Ile1102Thr, and His1086Leu to cluster together, except for MET wild type (with or without HGF), which clustered separately. This suggests similar gene expression programs in basal condition. Under HGF stimulation, the MET-activating mutations METex14Del, Ile1102Thr, and His1086Leu clustered together into a distinct group, which suggests similar gene expression patterns under ligand stimulation. Accordingly, Ile1102Thr and His1086Leu were found, under HGF stimulation, to differentially regulate 912 genes in common. This represents, respectively, ∼51% and ∼71% of their differentially regulated genes (Figure 6C). METex14Del, Ile1102Thr, and His1086Leu were found to differentially regulate 722 genes in common. Although all the activating mutations displayed similar gene-regulating effects, METex14Del appeared associated with a specific transcriptional program, with differential expression of 1037 genes not included among those highlighted for MET-activating mutations in the kinase N-lobe domain. To better understand the consequences of gene dysregulations due to the different MET receptor mutations in tumorigenesis, we performed associated gene enrichment analysis. Gene enrichment analysis by overrepresentation revealed several molecular processes and molecular functions common to the METex14Del and Ile1102Thr and His1086Leu mutations, including regulation of extracellular structure and matrix organization, locomotion, and cell migration (Figure 6D and Supplementary Figure S3A). This is consistent with the increase in wound healing observed with these variants. The regulated genes concerned include, for example, matrix metalloproteases (MMP1, 3, 9,10 and 13) and integrins (ITGB6 and ITGA2). Individual analyses confirmed that in response to HGF, METex14Del, Ile1102Thr, or His1086Leu induced increase in the expression of the matrix metalloproteases (MMP1, MMP10 and MMP13) (Supplementary Figure S3A, B and C). The ReactomePA database (reactome pathway database) revealed, in addition, that the METex14Del variant can specifically regulate genes involved in the NF-κB pathway, in cell cycle progression (through p53), and in proteasomal degradation (Supplementary Figure S3B). Individual analyses confirmed that in response to HGF, METex14Del induced strongrer upregulation of CDKN2B, involved in cell cycle regulation[64] and CD44, involved in p53 regulation[65], comparer to Ile1102Thr, or His1086Leu variants (Supplementary Figure S3D and E). Thus, HGF stimulation of either METex14Del, Ile1102Thr, or His1086Leu appears to trigger a same transcriptional program consistent with cell mobility and invasion, although additional genes appear to be regulated by METex14Del.

**Figure 6:**
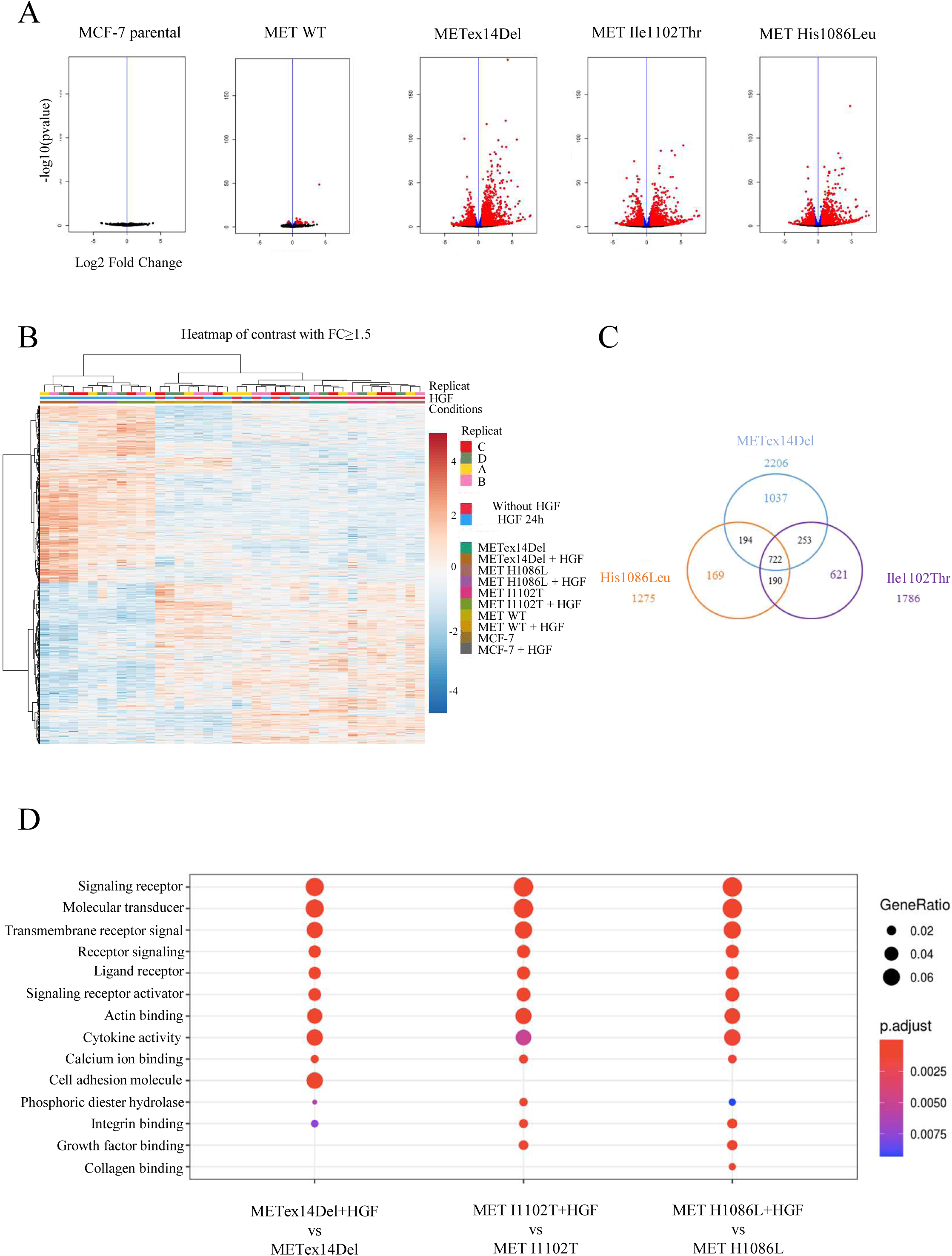
Transcriptional programs induced by the mutated forms of MET. Gene expression was determined by 3’RNA Seq on total RNA extracts of MCF-7 cells expressing either wild type (WT) MET, METex14Del, MET Ile1102Thr or MET His1086Leu, treated or not for 24 h with 30 ng/ml HGF in serum-free medium (n=4). (A) Volcano plot representation of differential expression analysis of genes in MCF-7 cells treated or not with HGF. Red and blue points mark, respectively, the genes with significantly increased or decreased expression (in blue padj<= 0.05, in red FC=> 1.5 & padj<= 0.05). (B) Heatmap of relative gene expression significantly upregulated (red) or downregulated (blue) in MCF-7 cells treated or not with HGF (fold change > 1.5 and p-value < 0.05; four replicates noted A,B,C,D were performed). (C) Venn diagram of genes significantly regulated in response to HGF stimulation in MCF-7 METex14Del, MET Ile1102Thr and MET His1086Leu cells (fold change >=1.5; ajusted p-value < 0.05). (D) Dot plot of gene ontology (GO) ‘molecular function’ enrichment among genes showing significant differential expression in MCF-7 cells according to whether they were stimulated or not with HGF (p.value adj <0.05 and fold change > 1.5).

### The I1102T substitution can induce experimental tumor growth

To investigate the ability of the MET-activating mutation to promote tumor growth, MET variants were stably transfected in NIH3T3, a cell line in which experimental tumor growth can be dependent on MET activity [66]. We obtained sufficiently high level of MET only for MET variants Met1268Thr and Ile1102Thr (Supplementary Figure S4), which were then xenografted into immunodeficient SCID mice in comparison with parental NIH3T3 cells. The mice injected with parental NIH3T3 cells did not develop any tumor. In contrast, all the mice injected with Met1268Thr cells developed tumors with a mean volume of 832 mm^3^. Mice injected with Ile1102Thr cells developed tumors (6 among 8), with a mean volume of 228 mm^3^ (Figure 7A). Expression of MET from tumor slides were assessed by immunofluorescence (Figure 7B). It is worth noticing that mice presenting tumors but sacrificed before the end point were not included in the figure. Our results thus demonstrate that Ile1102Thr expression can promote tumor growth, albeit less effectively than Met1268Thr expression. This tallies with the results of in vitro focus-forming assays (Figure 2). It suggests that Ile1102Thr can promote tumor formation in the absence of HGF stimulation and that its tumor-promoting capacity could be increased under HGF stimulation.

**Figure 7:**
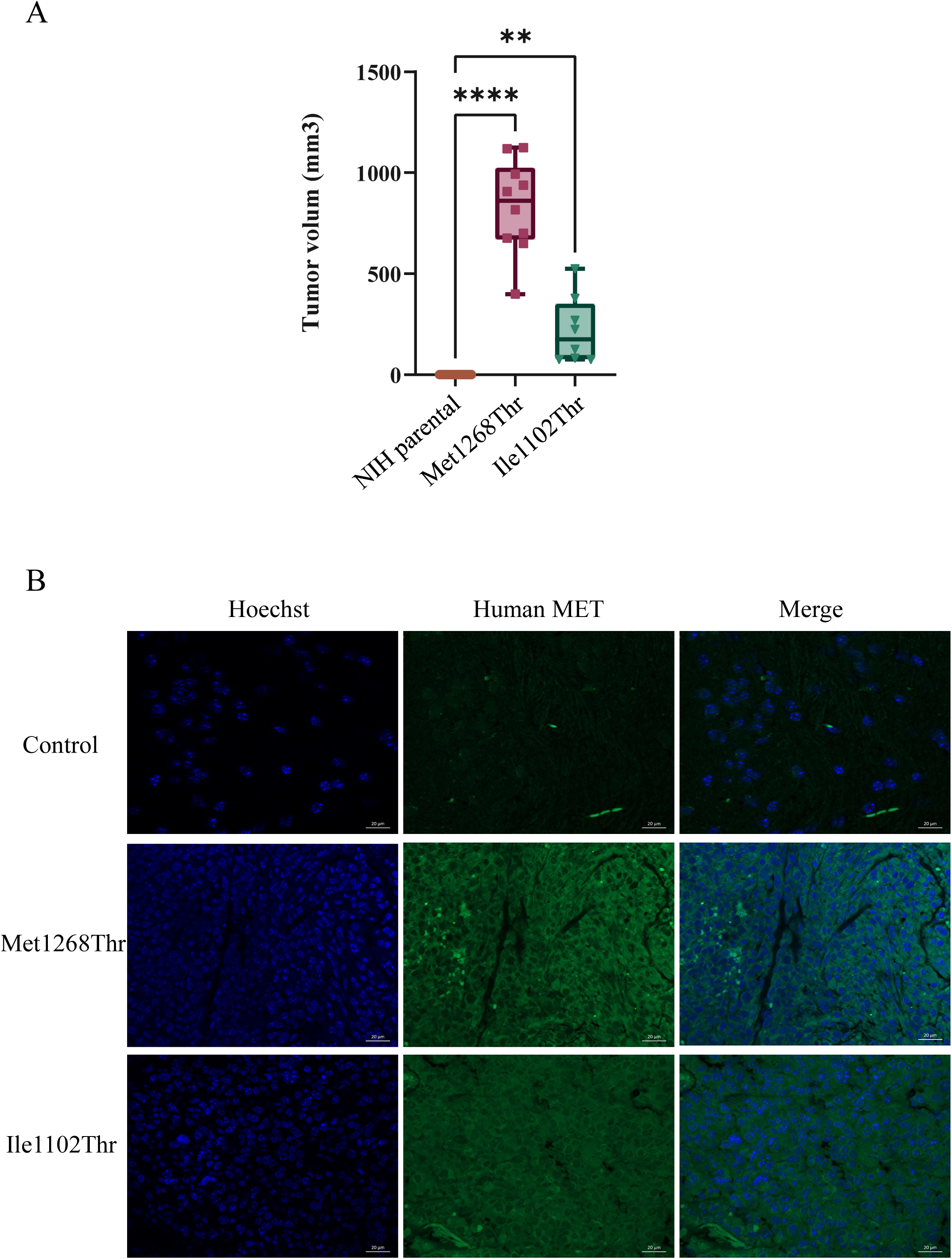
Experimental tumor growth of NIH3T3 cells expressing MET variants. In vivo tumor xenograft experiments were performed with SCID mice. (A) Scatter plots of the tumor volumes (mm3) at the end point in SCID mice xenografted with NIH3T3 parental cells, NIH3T3 cells stably expressing MET Ile1102Thr (8 measured tumors) or MET Met1268Thr (10 measured tumors). Error bars indicate the SEM; one way Anova test with: ** p-value <0.0021; **** p-value <0.0001.). (B) MET expression was analyzed by immunofluorescence in tumor slides of cells expressing MET M1268T and MET I1102T and adjacent murine tissue (control). Representative pictures of Hoechst staining, human MET staining and merge are shown. Scale bars represent 20 μm.

### The I1102T substitution may play a supportive role in ATP stabilization, whereas H1086L interrupts an ⍺JM-helix-⍺C-helix interface salt bridge

To investigate how the I1102T and H1086L substitutions act structurally as activating mutations, we mapped these residues on the human MET kinase domain crystal structure. Residue I1102T was mapped on the structure of the ATP-bound MET kinase domain (PDB 3DKC) (Figure 8A). The β1-β3 strands, P-loop, and hinge support the catalytic site where ATP binds. In the β1-β3 strands, H1112 and H1124 display polar interactions. I1102 is a residue in the β1 strand, at the entrance of the catalytic site (Figure 8B). The I1102T substitution is predicted to create a salt bridge interaction with the H1112 residue, which might modify the structure of the ATP binding site (Figure 8C).

**Figure 8:**
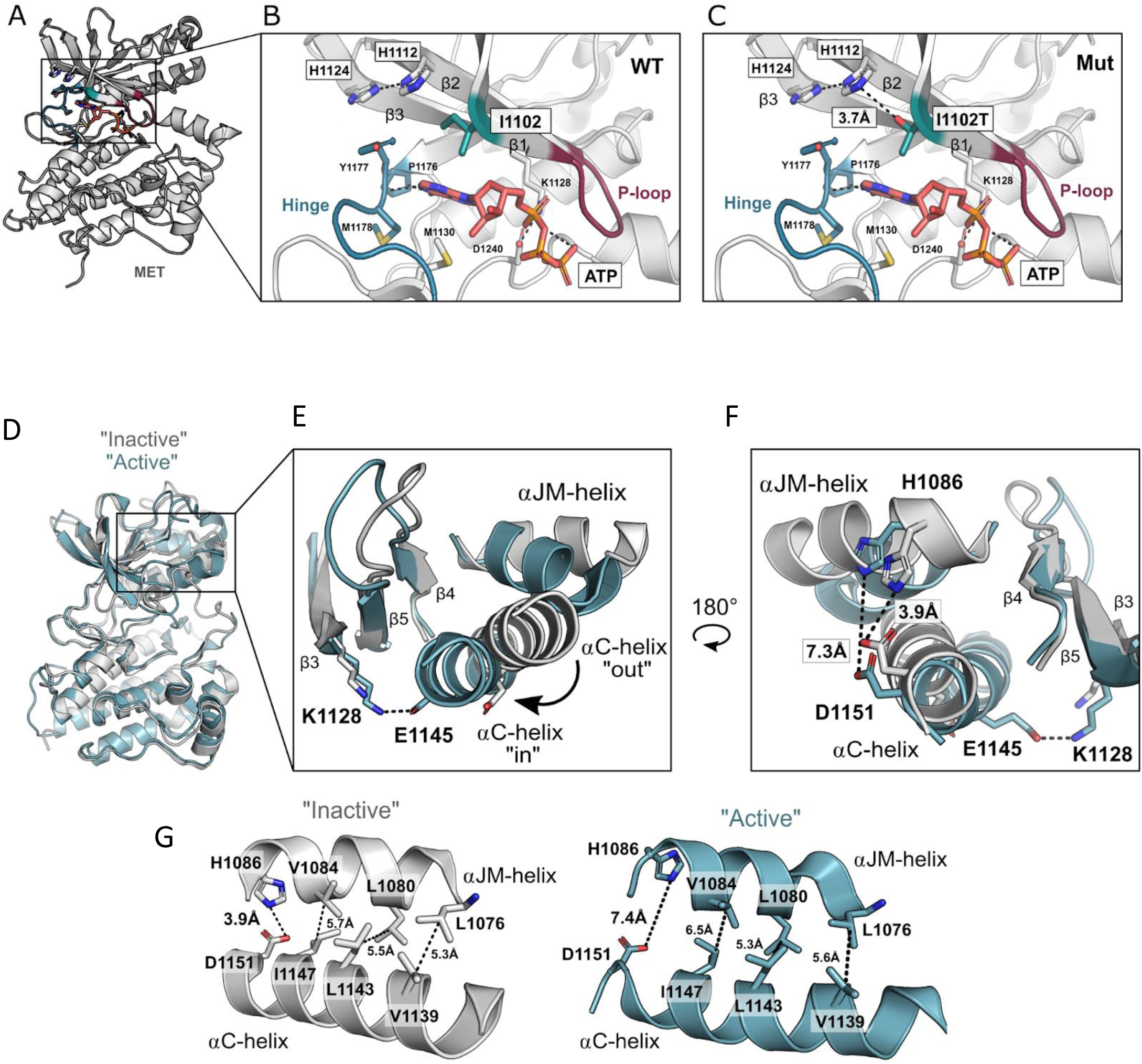
I1102T and H1086L MET mutations modeling in MET kinase structure. (A) Crystal structure of the ATP-bound human MET kinase domain (PDB 3DKC) analyzed through PyMOL. (B) ATP binding site and catalytic site motifs are highlighted in the kinase domain. The β1-β3 strands, P-loop (magenta), and hinge (teal) support the catalytic site, where ATP (orange) binds. Binding is stabilized by backbone interactions with hinge residues (P1176, M1178) and interactions with the conserved D1240 of the DFG motif and catalytic Lys (K1128). I1102 is a residue in the β1 strand of the N-lobe, at the entrance of the catalytic site. In the β1-β3 strands, H1112 and H1124 display polar interactions. (C) Representative “inactive” (gray, PDB 2G15) and “active” (blue, PDB 3R7O) human MET kinase domain crystal structures superimposed and analyzed through PyMOL. (D) Zoom-in view comparing how the ⍺JM-helix, ⍺C-helix, and β3-β5 motifs are siruated in an “inactive” and an “active” conformation: a conserved salt bridge between the β3 K1128 and ⍺C-helix E1145 is present in an “active” ⍺C-helix “in” conformation, and lost in an “inactive” ⍺C-helix “out” conformation. (E) One-hundred-and-eighty-degree rotation of the ⍺JM-helix, ⍺C-helix, and β3-β5 motifs between the “active” and “inactive” conformations. A salt bridge between the ⍺JM-helix H1086 and ⍺C-helix D1151 is present in an “inactive” ⍺C-helix “out” conformation and lost in the “active” ⍺C-helix “in” conformation. (F, G) Side-by-side view of the ⍺JM-helix-⍺C-helix interface in the “inactive” (gray) and “active” (blue) conformations. The interface between the JM-helix and the ⍺C-helix is supported by hydrophobic interactions, with the exception of a polar patch between H1086 and D1151. These take part in a salt bridge in the “inactive” conformation. Hydrophobic interactions between the ⍺JM-helix and ⍺C-helix are maintained in the “active” conformation, but the H1086 and D1151 salt bridge is lost, the distance between these residues going from 3.4 Å in the “inactive” conformation to 7.4 Å in the “active” conformation.

H1086 was mapped on the apo human MET kinase domain crystal structure (2G15) and on a representative “active” structure (3R7O)[67,68] (Figure 8D). MET has a canonical kinase domain structure, and H1086 is located in a structurally conserved helix belonging to the juxtamembrane sequence (⍺JM-helix) and packing on top of the ⍺C-helix [68,69]. The ⍺JM-helix and ⍺C-helix form a sensitive interface that is maintained by hydrophobic interactions between residues V1084/ I1147, L1080/L1143, and L1076/V1139 (Figure 8E). Mutations that disrupt these hydrophobic interactions have a drastic effect on MET activity [68]. H1086 is at the very end of the ⍺JM-helix, and along with D1151 of the ⍺C-helix, it is the only polar region of this interface. In an “inactive” conformation, a salt bridge (3.9Å) is formed between H1086 and D1151, and this salt bridge is lost in the “active” conformation (Figure 8F and G). Concomitantly, in the active form a salt bridge is formed between residue E1145 of the ⍺JM-helix and residue K1128 of the adjacent b3 sheet. This leads us to hypothesize that loss of the H1086/D1151 salt bridge might be important for this highly sensitive interface and might place the MET kinase in a state of pre-activation.

To test conservation of the H1086-D1151 salt bridge and its correlation with conformational states, we generated a set of 94 available human MET kinase domain crystal structures, measured the distance between the H1086 nitrogen and the oxygen of the D1151 carboxyl group, and filtered the structures on the basis of the following criteria: wild-type sequence, salt bridge between H1086 and D1151, salt bridge between L1128 and E1145. Within the set, two distinct, mutually exclusive conformational groups were identified: four structures (1R1W, 2G15, 3F66, 4KNB) with an H1086-D1151 salt bridge and three (3Q6W, 3R7O, 4IWD) with an L1127-E1145 salt bridge. This suggests the existence of two distinct states, consistent with either an active or an inactive conformation of the MET kinase, according to the relative position of the ⍺JM-helix.

## Discussion

EGFR-activating mutations found in the kinase domain of this protein are characterized as causing ligand-independent activation due to a conformational change of the ATP-binding pocket. As a result, the mutated EGFR receptor and its downstream signaling are constitutively activated independently of ligand binding[70]. In contrast, the MET exon14 mutations found in NSCLC patients lead to a different activation pattern, since the resulting receptor still requires ligand-stimulated activation[36,37]. This is consistent with deletion, in METex14Del, of a negative regulatory domain. We demonstrate here that the novel mutations Ile1102Thr and His1086Leu (identified in HPRCC and affecting the N-lobe) cause, in the absence of HGF, only minor focus formation in NIH3T3 cell cultures and are unable to promote epithelial cell motility and downstream pathway activation, whereas HGF stimulation increases focus formation, promotes cell migration, and strongly activates the RAS-ERK and PI3K-AKT signaling pathways. Taken together, these results demonstrate that the novel MET variants mutated in the N-lobe of the kinase domain can be classified as ‘HGF-dependent’, like the variants resulting from MET exon 14 skipping.

In order to understand how these mutations allow HGF-dependent MET activation, they were modeled on the MET kinase structure. Interestingly, Ile1102Thr is located at the entrance of the catalytic site and could disrupt the ATP binding site. It is worth noting that mutation of the residue homologous to MET I1102 in ERK2, EGFR, BRAF, and HER2 causes resistance to kinase inhibitors[60]. This is not the case of the MET Ile1102Thr mutation, as concerns the tested MET TKIs. It would thus seem that although the mutation disrupts the ATP binding site, it does not perturb TKI anchoring. The His1086Leu mutation is located in the ⍺JM-helix and could disrupt the salt bridge present in the inactive form and absent from the active one. Although modeling provides information about the activating potential of variants, why HGF stimulation is still required to fully activate these MET variants remains unclear. We speculate that the identified mutations might not be able to fully activate the kinase, and that other conformational modifications might be required, such as opening of the activation loop, which could depend on HGF stimulation.

Although neither mutations causing MET exon 14 skipping nor the studied N-lobe substitutions render MET activation HGF-independent, the effects of these mutations are not altogether identical. For example, the results of our time course experiments confirm, as previously demonstrated, that MET exon 14 skipping induces sustained activation of downstream signaling[36,62,63,71–73], in contrast to the new N-lobe mutations. This is consistent with loss of the juxtamembrane domain, involved in receptor degradation, when exon 14 is skipped. Our transcriptional analysis further demonstrates that similar sets of genes, involved in cell motility and invasion, are induced in response to HGF in MET exon14 and N-lobe kinase mutants. Yet the corresponding gene expression profiles observed in this analysis overlap only partially: the MET exon14 variant appears to regulate more genes, notably involved in proliferation and proteasomal degradation, possibly as a consequence of sustained signaling pathway activation. A time course transcriptional analysis after HGF stimulation could provide additional information about the transcriptional program induced by these MET variants.

Since HGF expression by the tumor itself or by its microenvironment may be required for full activation of MET variants having an N-lobe mutation in HPRCC, HGF expression could be a valuable biomarker in this cancer. We have previously demonstrated that in NSCLC tumors harboring MET exon 14 mutations, HGF is expressed in about half of the tumor cells. This suggests the establishment of an autocrine loop leading to MET activation and tumorigenesis[36]. It would therefore be interesting to determine whether HPRCC tumors express HGF, which could further activate MET. It is worth noting that HPRCC tumors frequently display duplication of chromosome 7, carrying the HGF gene[29].

Lastly, we demonstrate here that the identified N-lobe MET variants remain sensitive to the type I and type II MET TKIs tested here. Although the Ile1102Thr mutation lies close to the ATP binding site and the homologous mutation in CDK6 results in resistance to ATP competitors[60], MET Ile1102Thr remains sensitive to MET TKIs. Therefore, patients harboring these novel N-lobe mutations are potentially eligible for MET TKI treatment. Since these mutations depend on HGF stimulation, targeted therapies preventing ligand/receptor binding, including antibodies against MET or HGF, might be used for these patients.

## Conclusion

MET variants in the N-lobe of the kinase domain, found in hereditary papillary renal cell carcinoma, still require ligand stimulation to promote cell transformation, in contrast to other RTK variants. This suggests that HGF expression in the tumor microenvironment is important for tumor growth in patient harboring such mutations. Their sensitivity to MET inhibitors opens the way for use of targeted therapies in these patients.

## Supporting information

Supplementary Table 1A Parental_HGF vs parental

Supplementary Table 1B MET_WT_HGF vs MET_WT

Supplementary Table 1C MET_Ex14_HGF vs MET_Ex14

Supplementary Table 1D MET_I1102T_HGF vs MET_I1102T

Supplementary Table 1E MET_H1086L_HGF vs MET_H1086L

## Conflict of interest

The authors declare that there is no conflict of interest regarding the publication of this article

## Acknowledgements

This work was supported by the CNRS, INSERM, the Pasteur Institute of Lille and the University of Lille, and by grants from the “Agence Nationale de la recherche”, “Cancéropôle Nord-Ouest”, the “Ligue Contre le Cancer, Comités Nord et Aisne” the “SIRIC ONCOLille” and the “Institut National du Cancer”. Canther laboratory is part of ONCOLille institute. This work is supported by a grant from Contrat de Plan Etat-Région CPER Cancer 2015-2020. The JSF work is supported by NIH CA239604.

## Data Availability statement

The data that support the findings of this study are openly available in The Sequence Read Archive (SRA), reference number BioProject ID : PRJNA1190835.

## Author Contribution Statement

CG and DT conceived and planned the experiments. CG, AV, MFe, ID, AL, RT performed experiments and processed the data. ABC, ER and CD contributed to patient samples collection and data analysis. JPM, CV and MFi performed transcriptomic and analyzed the data. GOE and JSF interpreted the crystal structures and proposed modeling. CG and DT wrote the manuscript with input from all authors. DT contributed to funding acquisition and directed the project.

## Supporting Information

**Supplementary figure S1:** Effect of an anti-HGF antibody treatment on wound healing induced by HGF on MCF-7 cells

**Supplementary figure S2:** Effect of MET tyrosine kinase inhibitors (TKI) on MET phosphorylation and downstream signaling pathways activation.

**Supplementary figure S3:** Gene enrichment analysis of MET variants under HGF stimulation.

**Supplementary figure S4:** Expression of selected genes from the transcriptomic analysis.

**Supplementary figure S5:** MET receptor expression determined by RT-qPCR in NIH3T3 cells stably transfected with MET variants.

**Supplementary Table 1A:** Differential gene expressions in parental cells stimulated or not by HGF

**Supplementary Table 1B:** Differential gene expressions in cells expressing MET WT stimulated or not by HGF

**Supplementary Table 1C:** Differential gene expressions in cells expressing METex14Del stimulated or not by HGF

**Supplementary Table 1D:** Differential gene expressions in cells expressing MET I1102T stimulated or not by HGF

**Supplementary Table 1E:** Differential gene expressions in cells expressing MET H1086L stimulated or not by HGF

## Notes

### Competing Interest Statement

The authors have declared no competing interest.

### Summary of Updates

Minor modifications in text to impove understanding.

