## Supplementary Figure S1 for "MET variants with activating N-lobe mutations identified in hereditary papillary renal cell carcinomas still require ligand stimulation"

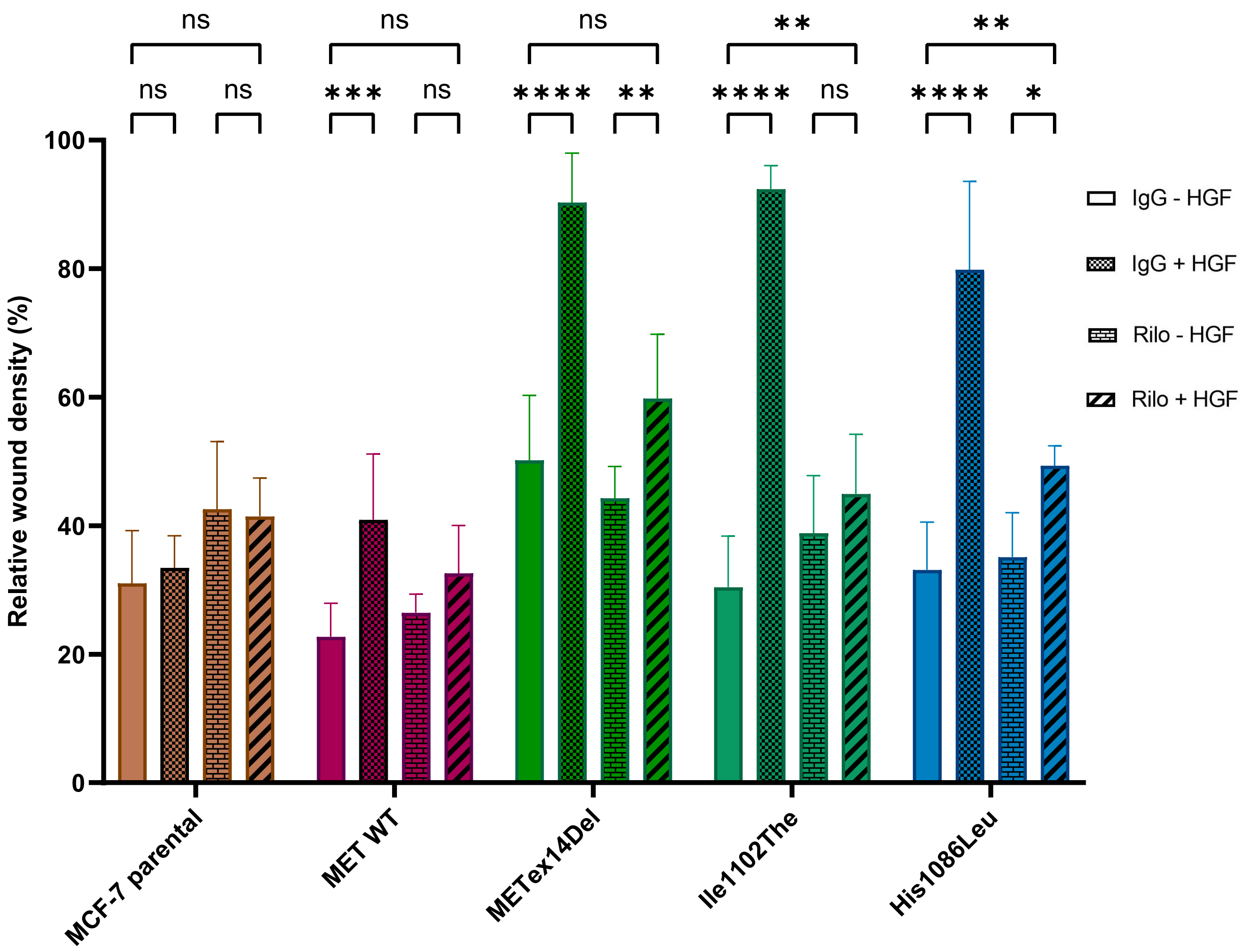


**Supplementary Figure S1**: **Effect of an anti-HGF antibody treatment on wound healing induced by HGF on MCF-7 cells**. Cells were seeded at 15,000 cells per well in a 96-well plate. 48h later, mitomycin C (10µg/mL) was added for 2h to prevent proliferation. A scratch wound was then performed and cells were stimulated or not with HGF at 30 ng/mL and Rilotumumab (Rilo) or control IgG (IgG) was added at 10µg/ml. Data are expressed as relative wound density 96h after HGF stimulation.*n*=4; mean ± SEM ; representative of three independent experiments. Statistical analysis by two-way ANOVA p<0.1234 (ns = not significant), p<0.0332 (*), p<0.0021 (**), p<0.0002 (***), p<0.0001 (****).
