## Supplementary Figure S2 for "MET variants with activating N-lobe mutations identified in hereditary papillary renal cell carcinomas still require ligand stimulation"

A

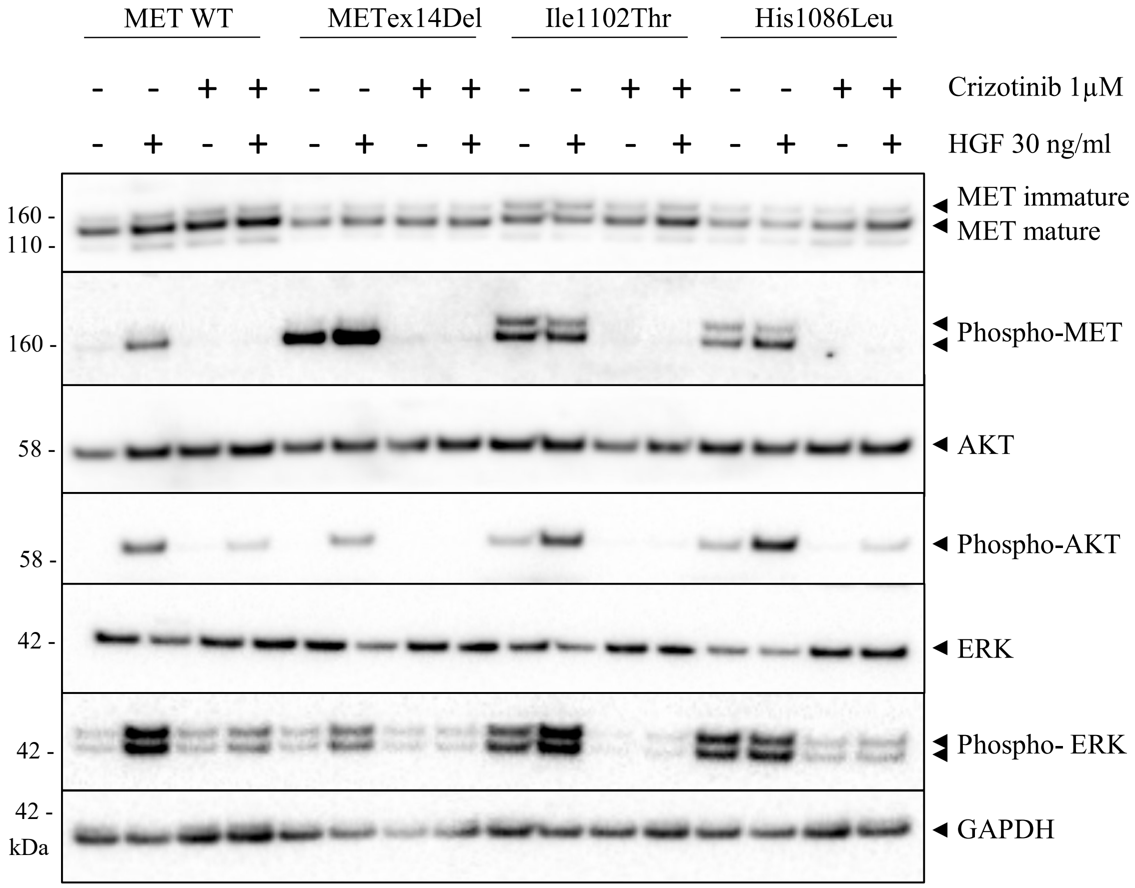
MET WT
METex14Del
Ile1102Thr
His1086Leu
Crizotinib 1µM
HGF 30 ng/ml
MET immature
MET mature
Phospho-MET
AKT
Phospho-AKT
ERK
Phospho- ERK
GAPDH
160 -
110 -
58 -
42 -
kDa

B

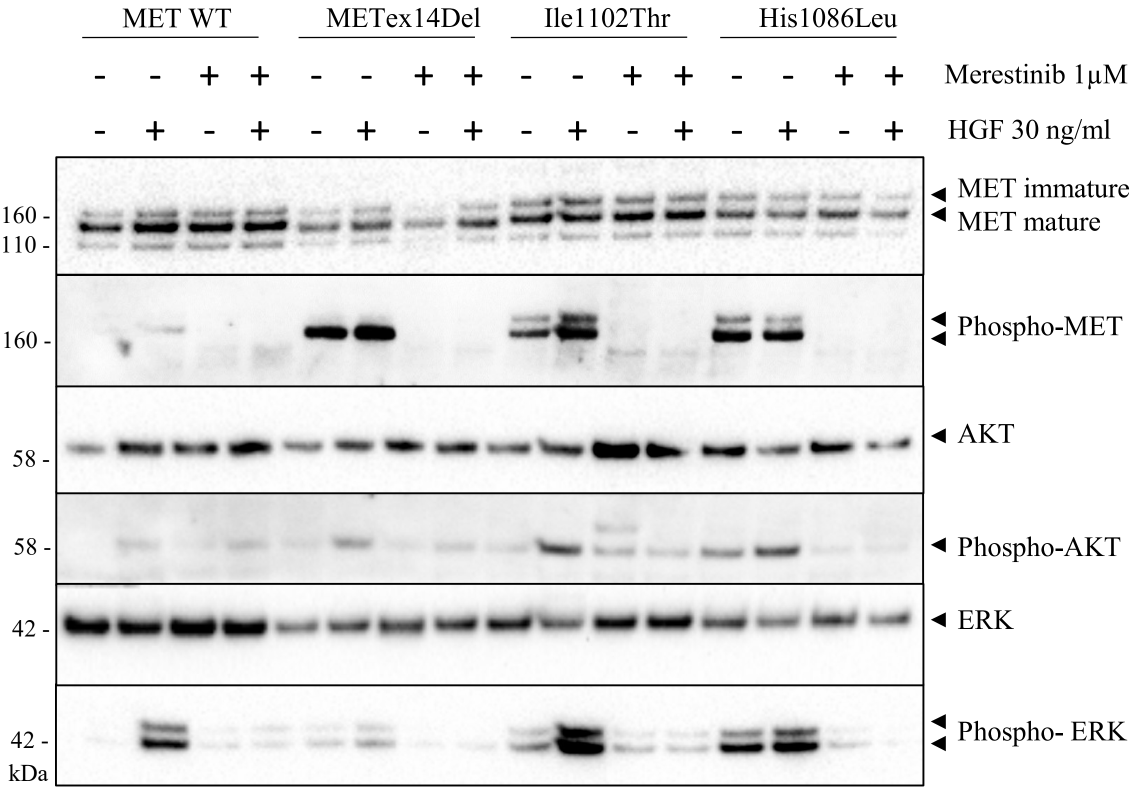
MET WT
METex14Del
Ile1102Thr
His1086Leu
Merestinib 1µM
HGF 30 ng/ml
MET immature
MET mature
Phospho-MET
AKT
Phospho-AKT
ERK
Phospho- ERK
160 -
110 -
58 -
42 -
kDa

C

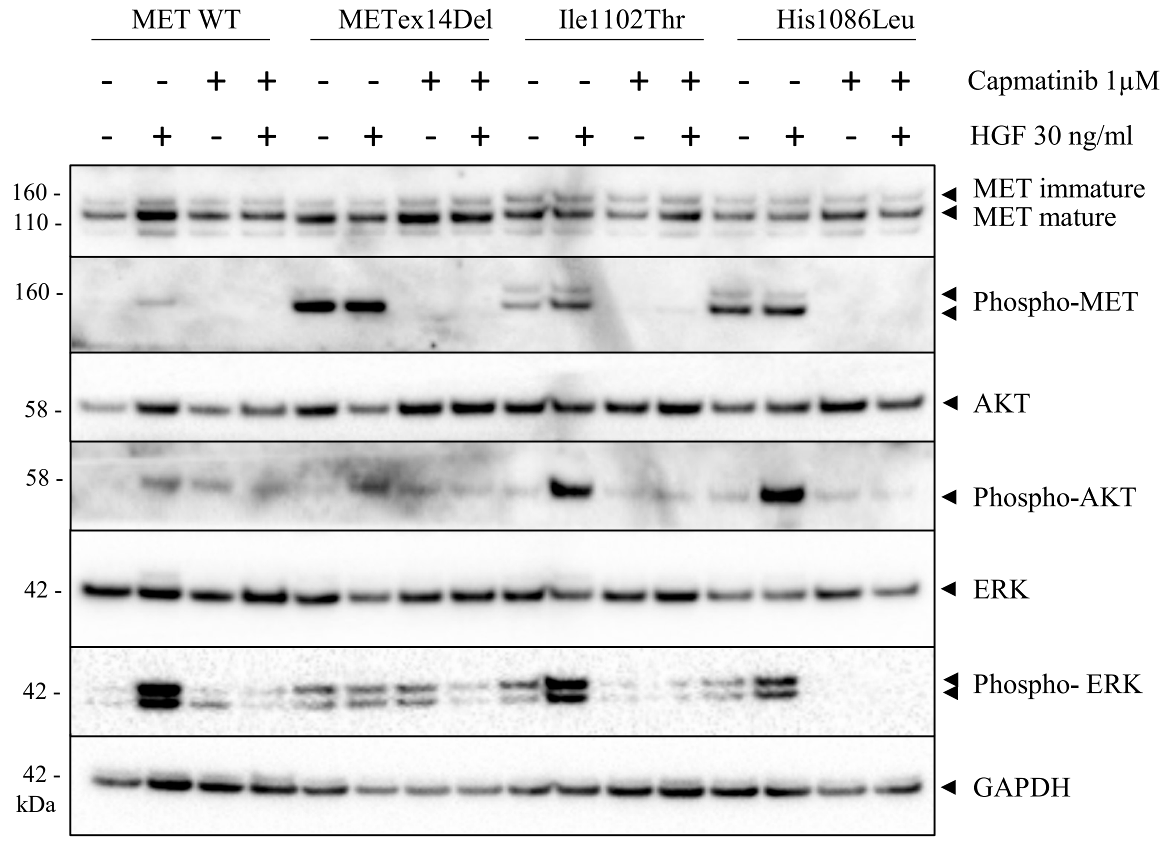


**Supplementary Figure S2**: **Effect of MET tyrosine kinase inhibitors (TKI) on MET phosphorylation and downstream signaling pathways activation.** MCF-7 cells expressing wild-type or mutated MET as indicated were incubated overnight in serum-free medium and then treated 1.5h or not by MET TKI (crizotinib (A), merestinib (B), or capmatinib (C) at 1μM) and stimulated or not for 30min with 30ng/ml HGF prior to cell lysis. Levels and phosphorylation of MET, AKT, and ERK, were determined by western blotting with the indicated antibodies. GAPDH was used as loading control.
