## Supplementary Figure S3 for "MET variants with activating N-lobe mutations identified in hereditary papillary renal cell carcinomas still require ligand stimulation"

A

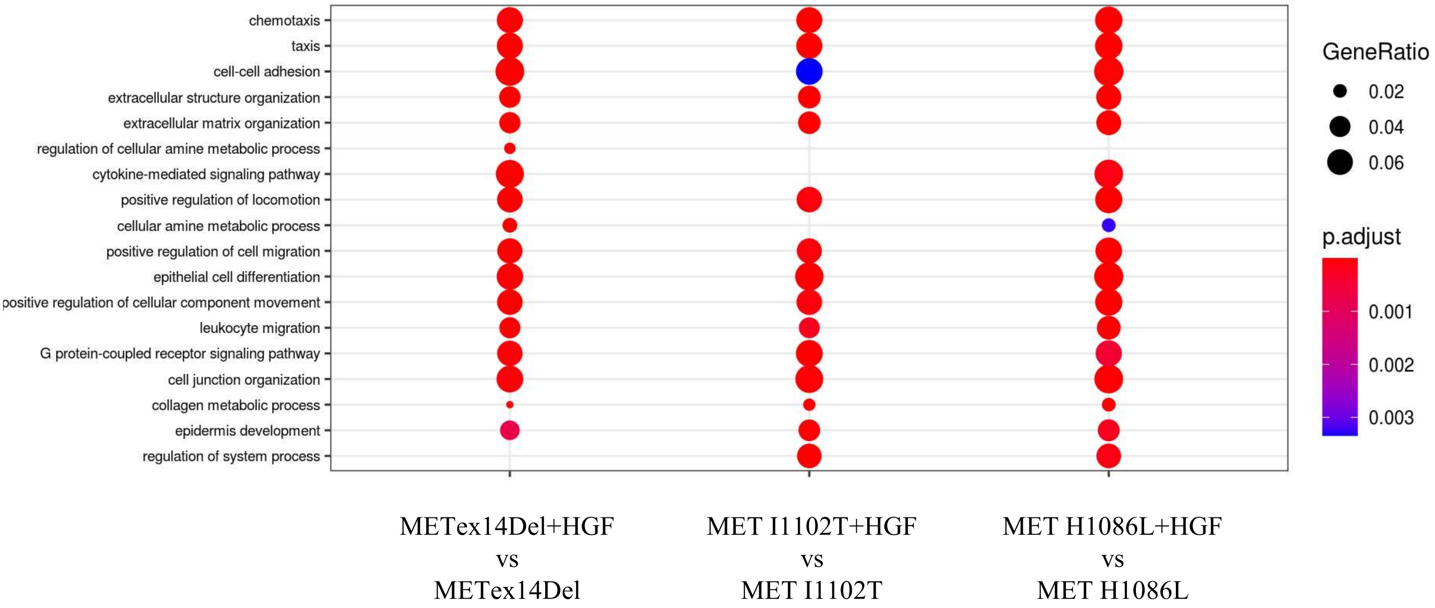


B

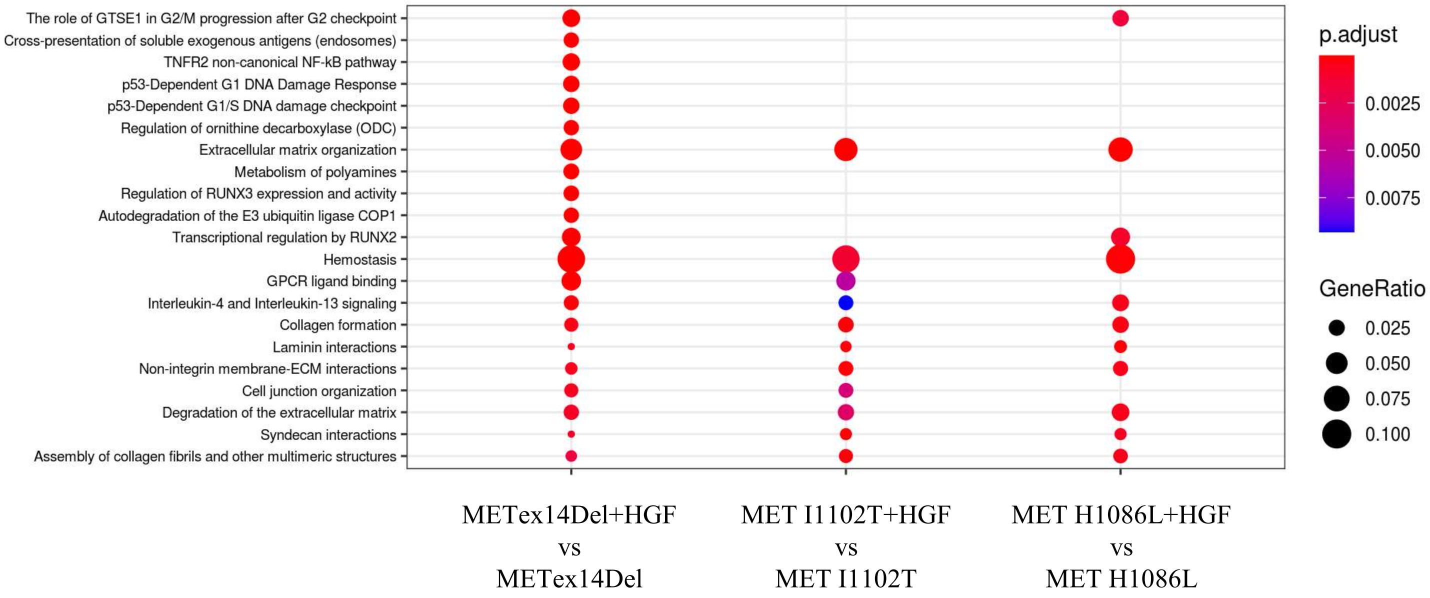


**Supplementary Figure S3**: **Gene enrichment analysis of MET variants under HGF stimulation.** (A) Dot plot of GO enrichment in 'Biological process' (A) and 'ReactomePA' (B) annotations among genes showing significant differential expression in MCF-7 cells according to whether they were stimulated or not with HGF (p.value adj <0.05 and fold change > 1.5). Dot size represents the gene ratio. Color scale represents the P-value (adjusted).
