## Supplementary Figure S4 for "MET variants with activating N-lobe mutations identified in hereditary papillary renal cell carcinomas still require ligand stimulation"

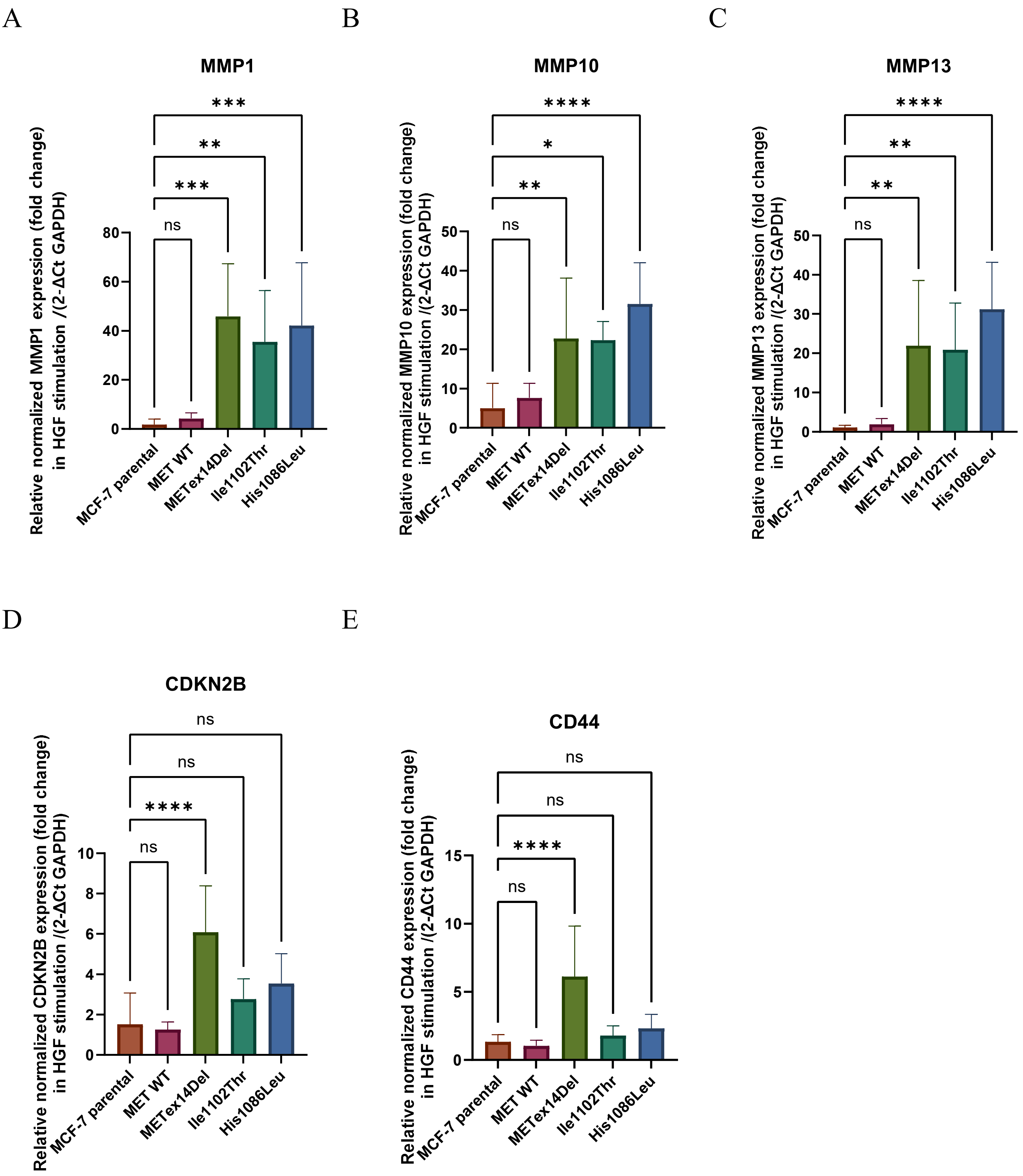


**Supplementary Figure S4**: **Expression of selected genes from the transcriptomic analysis.** Relative gene expressions to unstimulated conditions of MMP1, MMP10, MMP13, CDKN2A and CD44 determined by RT-qPCR in MCF-7 cells treated or not for 24 h with 30 ng/ml HGF in serum-free medium. * p-value <0.0332; ** p-value <0.0021; *** p-value <0.0002; **** p-value <0.0001.
