## Supplementary Figure S5 for "MET variants with activating N-lobe mutations identified in hereditary papillary renal cell carcinomas still require ligand stimulation"

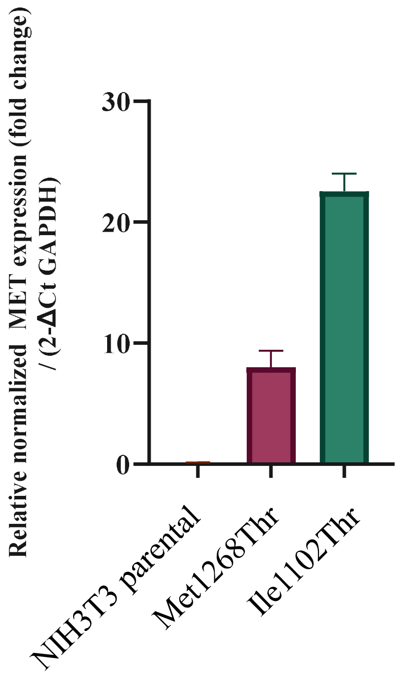


**Supplementary Figure S5: MET receptor expression determined by RT-qPCR in NIH3T3 cells stably transfected with MET variants.** Level of MET expression was determined by RT-qPCR in parental NIH3T3 and cells stably transfected with MET Met1268Thr and MET Ile1102Thr .
